# Ramified rolling circle amplification for efficient and flexible synthesis of nucleosomal DNA sequences

**DOI:** 10.1101/676528

**Authors:** Clara L. van Emmerik, Ivana Gachulincova, Vincenzo R. Lobbia, Mark A. Daniëls, Hans A. Heus, Abdenour Soufi, Frank H.T. Nelissen, Hugo van Ingen

## Abstract

Nucleosomes are a crucial platform for the recruitment and assembly of protein complexes that process the DNA. Mechanistic and structural *in vitro* studies typically rely on recombinant nucleosomes that are reconstituted using artificial, strong-positioning DNA sequences. To facilitate such studies on native, genomic nucleosomes, there is a need for methods to produce any desired DNA sequence in an efficient manner. The current methods either do not offer much flexibility in choice of sequence or are less efficient in yield and labor. Here, we show that using ramified rolling circle amplification (RCA) milligram amounts of a genomic nucleosomal DNA fragment can be produced in a scalable, one-pot reaction overnight. The ramified RCA reaction is more efficient than the existing methods, is flexible in DNA sequence and shows a 10-fold increase in yield compared to PCR, rivalling the production using plasmids. We demonstrate the method by producing the genomic DNA from the human LIN28B locus and show that it forms functional nucleosomes capable of binding pioneer transcription factor Oct4.

## INTRODUCTION

Nucleosomes are the repeating unit of chromatin, protecting genome integrity and regulating DNA-templated processes like transcription, replication and repair. This makes them a crucial scaffold for protein binding and therefore a highly attractive target for biochemical and structural studies. Nucleosomes are usually reconstituted *in vitro* from the individual histones H2A, H2B, H3 and H4 and a ~150 bp DNA sequence. To obtain stable, homogeneous samples, so-called strong-positioning sequences, such as the Widom 601 sequence (1) or the human a satellite repeat (2), are commonly used to ensure uniform positioning of the nucleosomes on the DNA. Although these sequences are widely used in the field, there is a concern that the resulting nucleosomes are more stable than native ones and therefore do not accurately reflect the situation *in vivo*. Therefore, there is an increasing interest in the study of nucleosome dynamics and interactions reconstituted from alternative or genomic DNA sequences. Since structural studies usually require milligram quantities of nucleosomes at high concentration, there is a demand for efficient procedures to obtain nucleosomal DNA with any sequence of choice.

At present, two methods are regularly used to produce nucleosomal DNA in milligram quantities. Firstly, the DNA can be produced from a plasmid containing multiple repeats of the desired sequence (3). The plasmid is transformed into *E. coli* and amplified by culturing the bacteria. The plasmid is then isolated and digested into the individual repeats, and the plasmid backbone is removed from the product sequence by ion exchange chromatography. In our lab, we get an average yield of 20 mg of product from 3 liters of culture using a 12-mer repeat of the 601 sequence; the Luger lab reported 46-61 mg from 6 liters of culture using a 24-mer repeat of the a satellite sequence. Alternatively, regular PCR amplification can be employed for nucleosomal DNA synthesis, starting from a plasmid containing a single copy of the desired sequence. This method is best used for small amounts, but can be scaled up, yielding 2-3 mg of pure product from 40 96-well plates in our lab. The plasmid method is usually preferred, as the yield is much higher. However, constructing the template plasmid containing multiple copies of the desired sequence can be challenging, forming a bottle neck when switching to different sequences. Although the single repeat plasmid used in the PCR method does allow sequence flexibility, the method is rather labor-intensive, and the yield is low.

A third option to produce DNA is based on rolling circle amplification (RCA). RCA requires a circular template and a polymerase that has a high processivity and strand displacement capacity, resulting in a long single-stranded product containing many complementary repeats of the template sequence. An advantage of RCA over PCR is that it is carried out in a one-step, isothermal reaction. RCA was originally developed for single-stranded amplification of so-called ‘padlock’ probes for specific DNA sequence detection (4–6). Since then, RCA has been used in a wide variety of applications, ranging from single molecule detection methods to the synthesis of DNA nanostructures and materials (for a review see (7)). For large-scale synthesis of single-stranded DNA, the RCA protocol has been extended to include digestion of the long RCA product into monomers by ‘cutter hairpins’ (8) or annealing of a complementary digestion splint to form double-stranded restriction sites (9). This procedure has been used to produce ‘monoclonal’ single-stranded DNA oligonucleotides on a microgram scale (8) and singlestranded DNA aptamers on a multi-milligram scale (9).

RCA can be tuned to produce double-stranded DNA by the addition of a primer complementary to the product strand. This leads to ramification or (hyper)branching and an overall enhanced amplification factor of the reaction (10,11). This technique, also known as cascade RCA of exponential RCA, has been exploited in diagnostic and biosensing assays as well as sequencing of single cell genomes to improve detection of low abundance nucleic acid targets (10,12).

Here, we show that rRCA is an efficient and flexible method for large-scale production of dsDNA, in particular for nucleosomal DNA synthesis. The protocol introduced here enables the production of milligram amounts of double-stranded DNA in a scalable, one-pot overnight reaction. By proper design of the template sequence, the long double-stranded product can be directly digested into monomers by a dedicated restriction enzyme. We detail the procedure to design and construct the circular template, to efficiently digest the rRCA product and purify the desired nucleosomal DNA. Our procedure yields 2 mg from a 12 mL reaction, a 10-fold increase in yield compared to the same volume of PCR reactions. The major advantage of the method is the flexibility in choice of DNA sequence. We demonstrate the method by producing genomic DNA from the LIN28B locus and showing that it forms functional nucleosomes capable of binding pioneer transcription factor Oct4.

## RESULTS

To illustrate the efficient production of any nucleosomal DNA sequences of choice by rRCA, we here focus on reconstituting nucleosomes using a sequence from a well-defined genomic locus in human fibroblasts. The LIN28B locus on chromosome 6 contains a well-positioned nucleosome that is the binding site for pioneer factors Oct4, Sox2, Klf4 and c-Myc (OSKM) (13) and is crucial in reprogramming and pluripotency (14,15). A 162 bp sequence from the LIN28B locus was successfully used for OSKM binding assays by Soufi *et al*. (16) and as such is an excellent model system to test the functional quality of the nucleosomal DNA generated by ramified RCA.

The approach is outlined in Figure 1. Briefly, a dsDNA sequence of choice is amplified from a storage vector, one of its strands is purified and circularized to yield the ssDNA template for RCA (Figure 1A). In ramified RCA, this template is amplified using two primers, a starting primer and a branching primer, and Phi29 DNA polymerase. This enzyme is frequently used for RCA because of its high processivity and its strand displacement capacity (17), high 3’-5’ exonuclease activity and low error rate (18,19). The starting primer anneals to the circular template and is elongated into a long, single-stranded repeat to which the branching primer can anneal. In regular RCA, only an equimolar amount of starting primer to circular template is needed, but in rRCA both starting and branching primer need to be in excess to template. Combined with strand displacement by Phi29, this results in the synthesis of a very long, branched, double-stranded repeat of the desired sequence. It should be noted that both starting and branching primer contribute to the ramification, as can be seen in Figure 1B. Subsequent digestion and purification yields the final nucleosomal DNA fragment.

**Figure 1.**
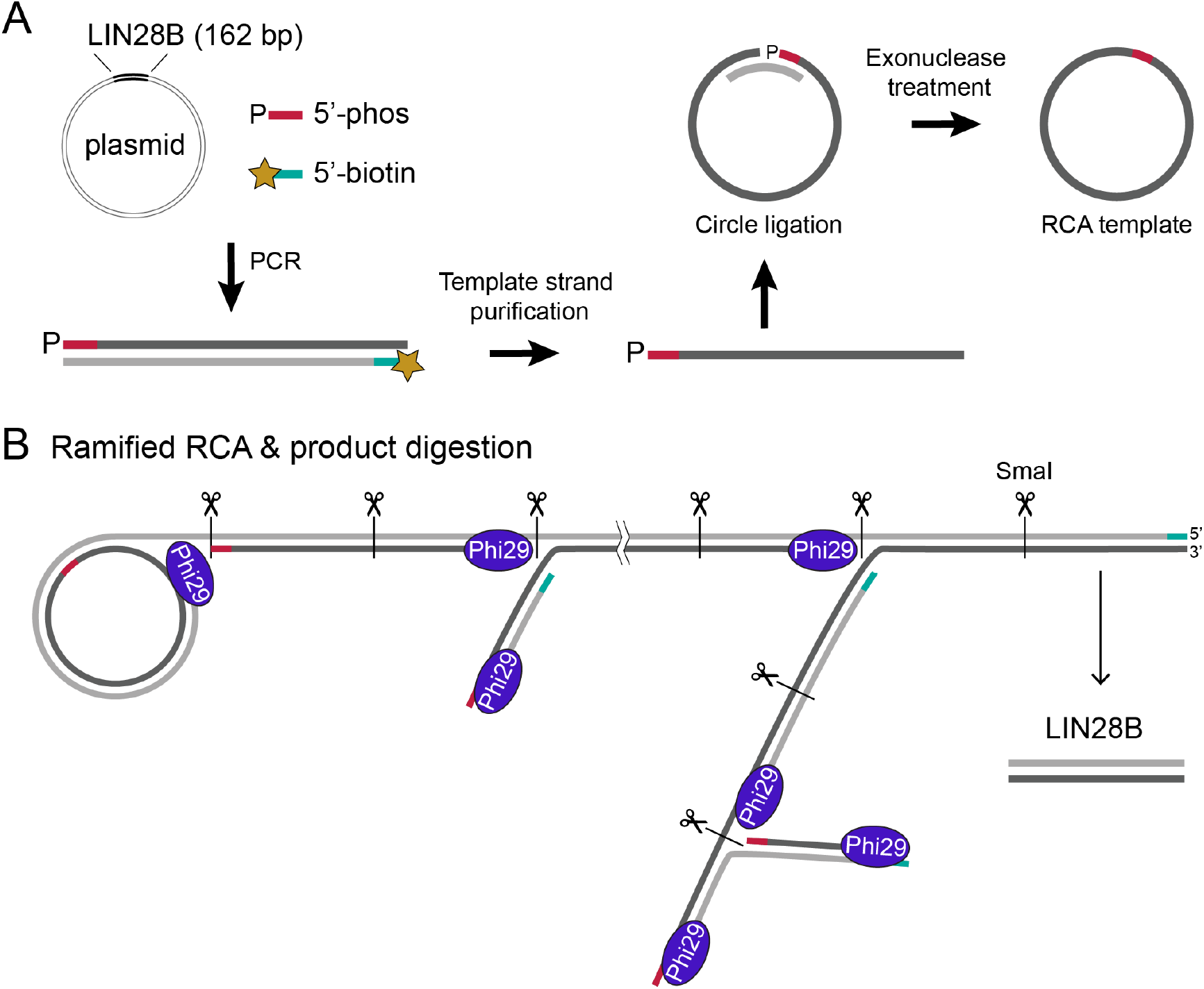
Schematic workflow of ramified RCA for nucleosomal DNA synthesis. (A) Preparation of circular templates, starting from a plasmid with the desired nucleosomal DNA sequence, a 5’-phosphorylated primer and a 5’-biotinylated primer. The template strand (dark gray) is purified by removing the biotin-labeled strand (light gray) using streptavidin beads. The template strand is circularized using a splint to anneal the 5’- and 3’-end together for ligation. (B) Ramified RCA starting from the circular template, two primers (pink and teal), and Phi29 polymerase produces a long, branched dsDNA product. Digestion is performed in the same reaction volume after addition of restriction enzyme SmaI (scissors) to release the double-stranded LIN28B product.

### Design and storage of the template

The DNA sequence of choice will typically need minor modification to allow efficient use in rRCA. First, the template sequence needs to be designed to encode a restriction site for a blunt-end nuclease in the rRCA product such that the final nucleosomal DNA fragment can be released. In the most straightforward design, the restriction site is formed from the ends of the linear template, and thus overlaps with the ligation site (see Fig. 1 and 2). Therefore, accurate ligation of these ends into the circular template is essential as mutations in the restriction site would consequently impair digestion of the rRCA product and significantly lower the final yield (see also below). Alternatively, the linear template can be designed in a way that the ligation site does not overlap with the restriction site, although care has to be taken not to introduce ligation mistakes that interfere elsewhere, e.g. with nucleosome positioning or protein binding. If ligation is found to be inaccurate, the ligation site can be separated from the template sequence by encoding a digestion site on both the 3’- and the 5’-end of the desired sequence, followed or preceded by a spacer sequence that contains the (half) ligation site such that it will be cut out during digestion of the rRCA product.

**Figure 2.**
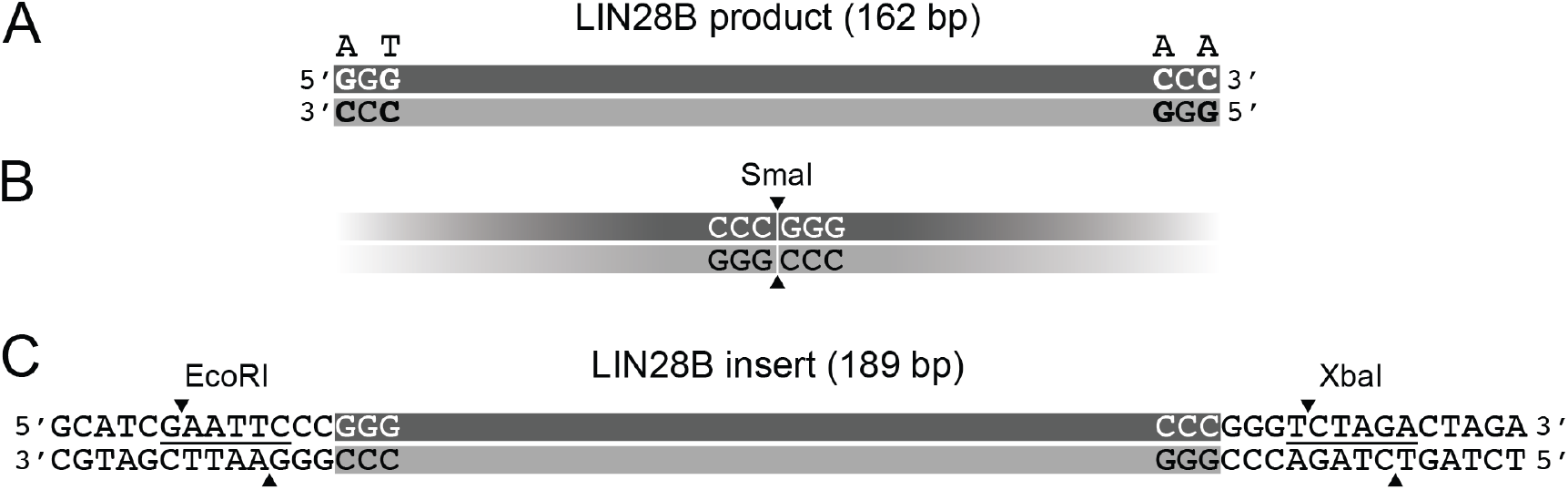
Template design of the LIN28B sequence for use in ramified RCA. (A) Four bases of the original LIN28B sequence were mutated (in bold, original base indicated on top) to create a SmaI restriction site in the multimeric RCA product (B). (C) The extended LIN28B insert contains two restriction sites (EcoRI and XbaI, underlined) to enable cloning into the pUC19 vector.

The choice of restriction enzyme can be guided by sequence of the terminal bases in the desired nucleosomal DNA. As long as these bases are not part of a functional protein binding site, they can likely be altered at will since nucleosome positioning is encoded within the central 120bp of the nucleosomal DNA. For LIN28B, we changed 4 residues at the termini of the original sequence to construct the recognition site for blunt-end cutter SmaI (CCCGGG), the first half of which (CCC) is at the 3’-end and the second half (GGG) at the 5’-end of the template sequence (Figure 2A,B). Notably, we found that Phi29 is highly active in CutSmart^®^ buffer when supplemented with 4 mM DTT. Thus, rRCA and digestion can be performed in the same reaction buffer, avoiding an additional buffer-exchange.

The RCA method starts from a circular, single-stranded DNA template. Single-stranded oligonucleotides up to 200 nucleotides are nowadays commercially available and could directly be circularized to use as an RCA template. However, the synthesis yield of such long oligonucleotides is usually limited to a few nanomoles and sequence inhomogeneity at the 5’-end can reduce ligation efficiency or result in incorrect ligation products. This not only lowers the amount of desired circular templates available but may also give difficulties with the digestion of the product when the restriction site lies in or near the ligation junction. To create a robust stock of template, we therefore decided to generate the ssDNA template from a synthetic dsDNA gene. Such dsDNA fragments can be synthesized with highest sequence accuracy and can furthermore be stored in a vector, allowing for easy, reliable and low-cost amplification in *E. coli*. The desired ssDNA template can then be derived from this material as described below.

The LIN28B sequence was cloned in a pUC19 vector by extending the desired sequence with flanking restriction sites (Figure 2C). Note that these extensions will not end up in the final nucleosomal DNA fragment. This sequence was ordered as a sequence-verified gBlock^®^ double-stranded gene fragment (IDT). The final cloned constructs were verified by sequencing. This simple and straightforward cloning procedure can easily be performed with any other desired sequence and provides in a robust template for the preparation of rRCA single-stranded circles.

### Preparation of the template circles

To obtain the ssDNA template from the storage plasmid, a PCR using two 5’-modified primers is required (Figure 1A). The primer producing the template strand is 5’-phosphorylated, as the phosphate-group is necessary for ligation to the 3’-end. In order to purify this strand, the primer producing the other strand is 5’-biotinylated. Notably, either strand of a dsDNA molecule of interest can be chosen as template, as both will yield the same double-stranded final RCA product. By binding the PCR product to streptavidin beads and eluting the phosphorylated template strand with sodium hydroxide, pure single stranded product can be obtained. As the streptavidin beads have high binding capacity, no PCR product purification is necessary to remove excess biotinylated primers before binding to the streptavidin beads. This procedure yielded pure ssDNA LIN28B fragment in a straightforward manner (Figure 3A).

**Figure 3.**
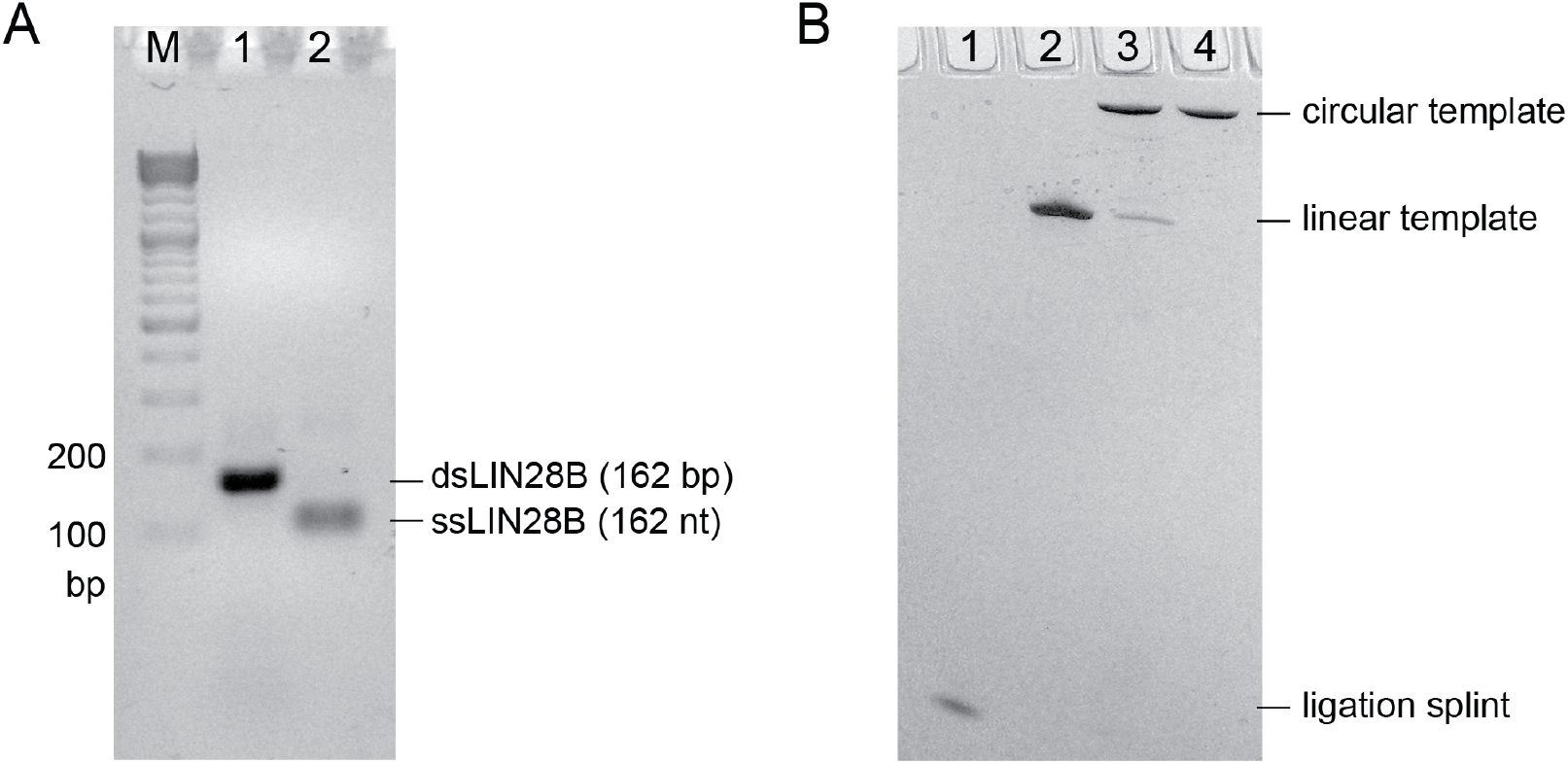
Template strand production and circularization. (A) Agarose gel (2.5%, EtBr staining) showing template strand purification: crude PCR product (lane 1) and purified LIN28B strand (lane 2). (B) Denaturing PAGE gel (12%) showing circularization and exonuclease digestion of the template strand: Ligation splint (lane 1), purified template strand (lane 2), ligated template circles, crude (lane 3) and after exonuclease clean-up (lane 4).

To circularize the linear template strand a short ssDNA fragment, the ligation splint, is needed to anneal both ends together. This creates a double-stranded region with a nick that will be ligated using a DNA ligase. Although T4 DNA ligase is a very efficient and commonly used enzyme for ligating nicks, it is also capable of ligating the termini if there is a 1 or 2 nucleotide gap present or if the nick contains mismatches (20–22). These gaps and mismatches could occur due to incorrect annealing of both ends on the ligation splint or by heterogeneity in the 5’-end of the primer used in PCR, commonly present in synthetic oligonucleotides. Since inaccurate ligation will cause mutations in the restriction site, special care needs to be taken to avoid this. We therefore used the thermostable Taq DNA ligase, which has a higher ligation accuracy and has no or little activity on gaps or mismatches at the ligation site (23). Taking advantage of its thermostability, the accuracy of splint annealing can be further increased by carrying out the ligation at elevated temperature. For LIN28B a 26nt ligation splint was used (see Table 1) in a ligation reaction with Taq DNA ligase at 45°C. The elevated temperature and thus more stringent annealing increased the amount of ligation efficiency, thus resulting in a homogenous pool of circular templates (Figure 3B).

**Table 1.**
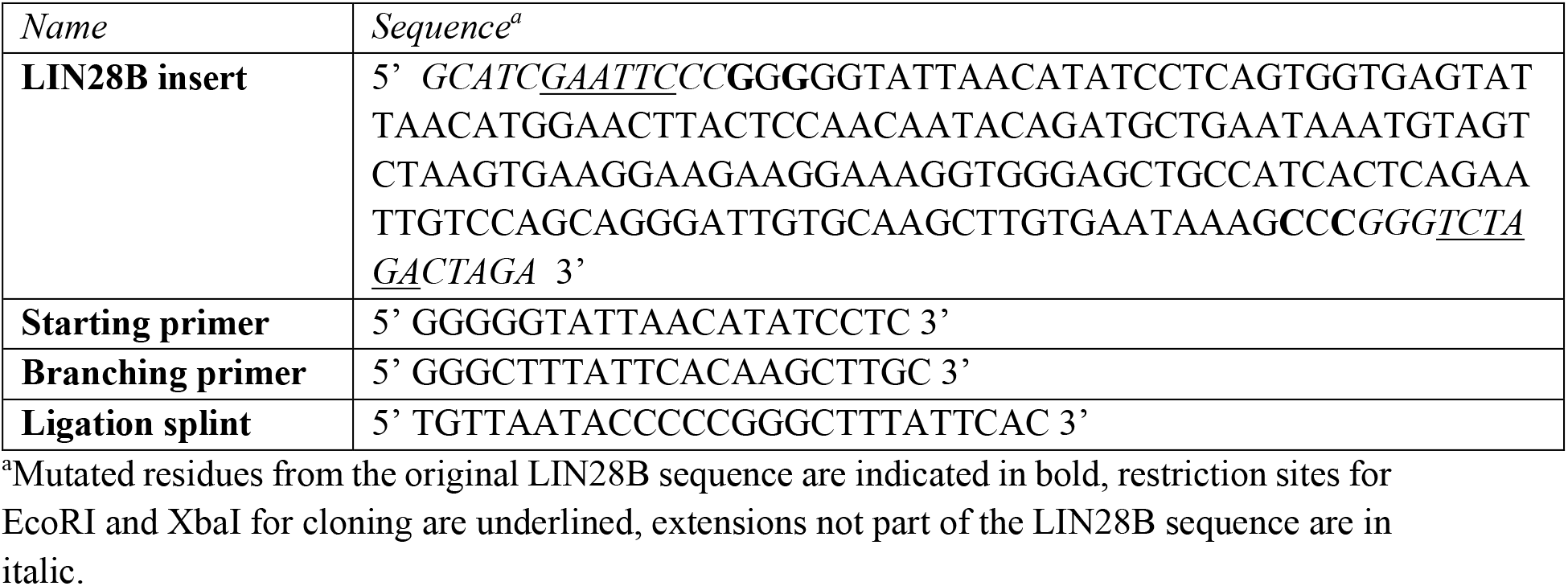
DNA sequences of the LIN28B gene fragment and oligonucleotides

After ligation, the reaction mix not only contains the circular template but also left-over splint and linear template. These could prime either on the circle or on the product during RCA, resulting in single stranded by-products. To avoid these contaminations of the product, all linear ssDNA fragments remaining in the ligation are digested with exonuclease I and III. Figure 3B shows this treatment results in pure circular templates. Sequencing analyses using the PCR primers confirmed that the purified circles contain the correct LIN28B sequence.

### Optimization of ramified RCA reaction conditions

Ramified RCA is performed as a one-pot reaction at constant temperature (30°C for Phi29 polymerase). The optimal reaction buffer will depend on both the polymerase and restriction enzyme used, as mentioned above. For rRCA, the yield depends on the degree of branching and thus on the amount of starting and branching primer. Both primers need to be added in a 1:1 ratio to avoid contamination with ssDNA. To maximize the yield, also the amount of dNTPs and Phi29 polymerase can be increased as these can become limiting. Finally, the purity of the circular template is essential to avoid by-products from remaining linear fragments (see above).

Initial rRCA reactions were performed using template circles that were purified after a standard 30-minute exonuclease treatment. While the templates appeared pure on gel-analysis, considerable ssDNA LIN28B product (and multimers) were generated after digestion, indicating presence of residues linear template or annealing splint (Figure 4A). By extending the exonuclease treatment three-fold, these by-products could be largely eliminated (Figure 4A). Unexpectedly, in negative controls carried out without starting and branching primers, RCA product formation was still observed when starting from these pure template circles (Figure 4B). This suggests that there are trace levels of DNA in the reaction mixture that can act as primer. These likely originate from the Phi29 polymerase stock, as co-purification of DNA is hard to avoid in preparations of this enzyme due to its high affinity for ssDNA and dsDNA (24).

**Figure 4.**
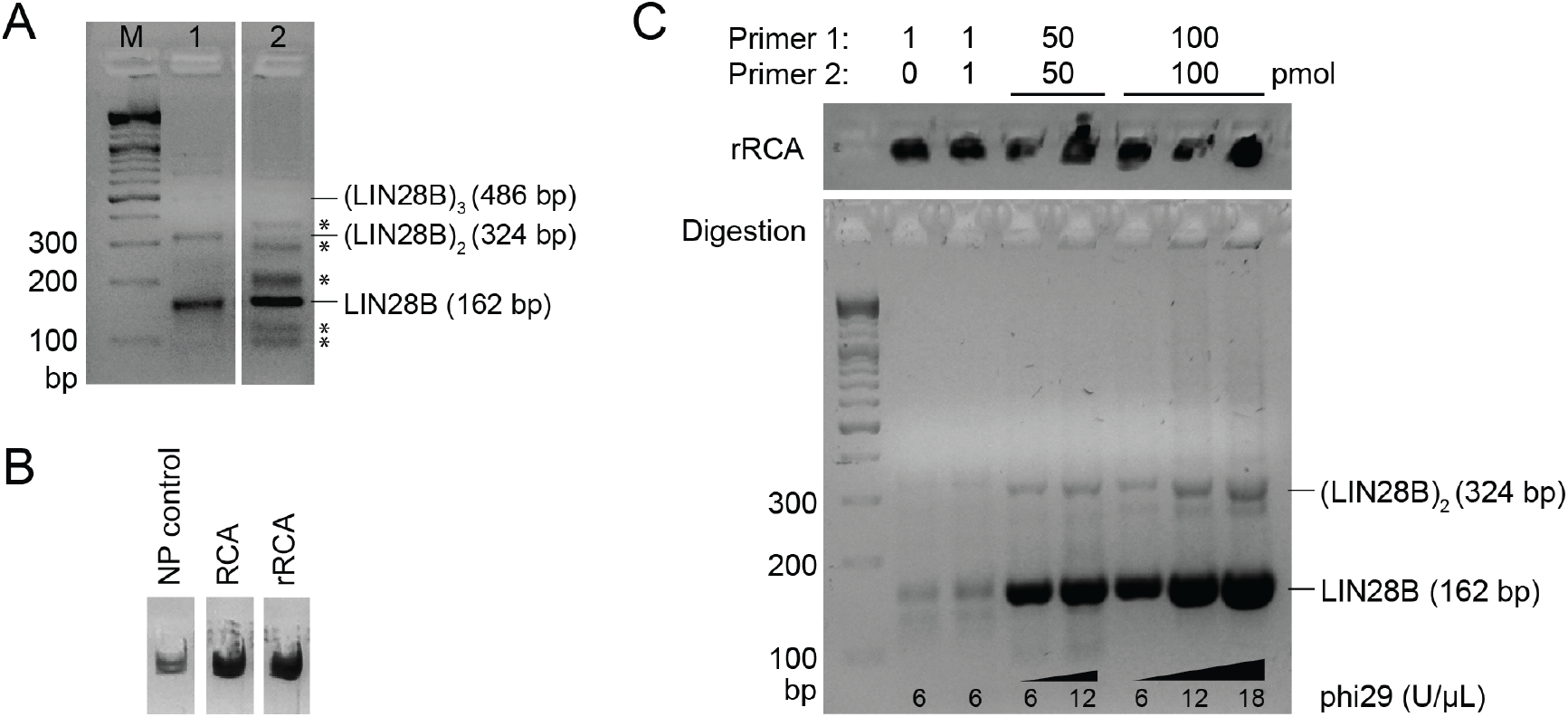
Optimization of ramified RCA reaction conditions. (A) Agarose gel (2.5%, EtBr staining) showing digested rRCA products from a reaction with a clean circular template (lane 1) and with a circular template that still contained impurities (lane 2). Single-stranded byproducts are indicated with an asterisk (*). (B) RCA products for ‘no primer’ (NP) control compared to product from RCA (1 pmol starting primer) and rRCA (50 pmol of both primers) on 0.8% agarose gel (EtBr staining). The circular template (1 pmol) used in this experiment was treated with exonuclease I and III for 1.5h. (C) Optimization of primer amounts for ramification and adjustment of Phi29 polymerase amount (2.5% agarose gel, EtBr staining). Top panel shows rRCA product, bottom panel shows SmaI-digested product. Before digestion, the rRCA yield is hard to compare, because of inconsistent pipetting due to high viscosity of the sample. Primer amounts are indicated above each lane, Phi29 concentrations are indicated below each lane.

To determine the optimal primer and Phi29 amounts, a series of 50 μL reactions were carried out using an excess of dNTPs (500 μM), and in presence of pyrophosphatase to avoid pyrophosphate-induced sequestering of the Mg^2+^ required for the polymerase. All reactions showed formation of a large RCA product that remains in the well of the agarose gel (see Figure 4B and Figure 4C, top panel). Since the product was difficult to pipet due to its high viscosity, the rRCA yield was assessed only after digestion by SmaI, resulting in LIN28B monomers of the expected size (162 bp) together with some dimers (324 bp) and multimers resulting from incomplete digestion (Figure 4C, bottom panel).

First, the non-ramified control reaction, using only starting primer in equimolar ratio to the circular template, generated double-stranded product of the same size as LIN28B monomers, while a long single-stranded product was expected (Figure 4C, lane 1). This may be caused by trace levels of DNA impurities as mentioned above, possibly in combination with aspecific priming of the starting primer or template-switching by Phi29 (25).

Second, addition of an equimolar amount of the branching primer showed double-stranded product formation with negligible increase in yield compared to the non-ramified reaction (Figure 4C, lane 2). Under these equimolar conditions, the branching primer can anneal to the product as soon as a full ssDNA LIN28B repeat is synthesized, resulting in only one or a few repeats of double-stranded product and no ramification, leaving the majority still singlestranded.

Reactions with either 50- or 100-fold molar excess of both primers, showed significant and progressively increasing yield of double-stranded product, indicating successful ramification (Figure 4C, lane 3 and 5). This confirms that effective branching only occurs at excess of both primers compared to the template. At these high primer concentrations, the amount of Phi29 polymerase becomes limiting, as seen by the increase in yield upon adding two- or three-fold more Phi29 at constant primer concentration (Figure 4C, lane 4, 6 and 7). Further increase of either primers or Phi29 may increase yield even more. Yet since very high excess polymerase is known to reduce polymerization efficiency and reaction volume can easily be scaled (see below), we choose to not explore even higher excess of this component.

In all cases, reaction time was limited to five hours. Since at this point the reaction mixtures turned highly viscous due to product formation, we presumed reaction progress would be significantly reduced and thus little benefit would be gained by extending the reaction for longer.

While these results highlight a sensitivity of Phi29 polymerase to the presence of trace levels of primers, large excess of starting and branching primer under ramified conditions will ensure correct priming and product formation. The results further demonstrate that maximum rRCA yield of dsDNA product is obtained at high molar excess of both primers in combination with elevated levels of Phi29 polymerase. These conditions are most likely independent of the DNA sequence being produced.

### Large-scale production, digestion & purification

For large-scale production of nucleosomal DNA, we take advantage of the fact that both rRCA and digestion are performed in a single reaction tube without the need of thermocycling.

This means the reaction can easily be scaled up by increasing the reaction volume and reaction components accordingly. For large-scale production of LIN28B DNA, we performed both a 2 mL and a 12 mL synthesis in a single tube. Since these large-scale rRCA reactions require larger amount of the polymerase, we use in-house produced Phi29 (see Materials and Methods) to reduce costs.

For LIN28B a 2 and 12 mL rRCA were performed under conditions determined above for 5 hours. In both cases, viscosity of the solution became very high at this point, which was quickly lowered after addition of the restriction enzyme SmaI due to the digestion of the rRCA product into 162bp LIN28B monomers. Restriction was almost complete after overnight incubation (Figure 5A). The remaining larger fragments were separated from the main product by anion exchange chromatography using a very shallow salt gradient, resulting in pure nucleosomal DNA (Figure 5B and C). We obtained 0.35 mg (3500 pmol) of purified product starting from 40 pmol of circular template in a 2 mL reaction volume, which is almost a 10-fold increase in yield compare to the same volume of regular PCR reactions. The 12 mL synthesis yielded 2.0 mg of pure LIN28B, demonstrating that the rRCA scales linearly with reaction volume.

**Figure 5.**
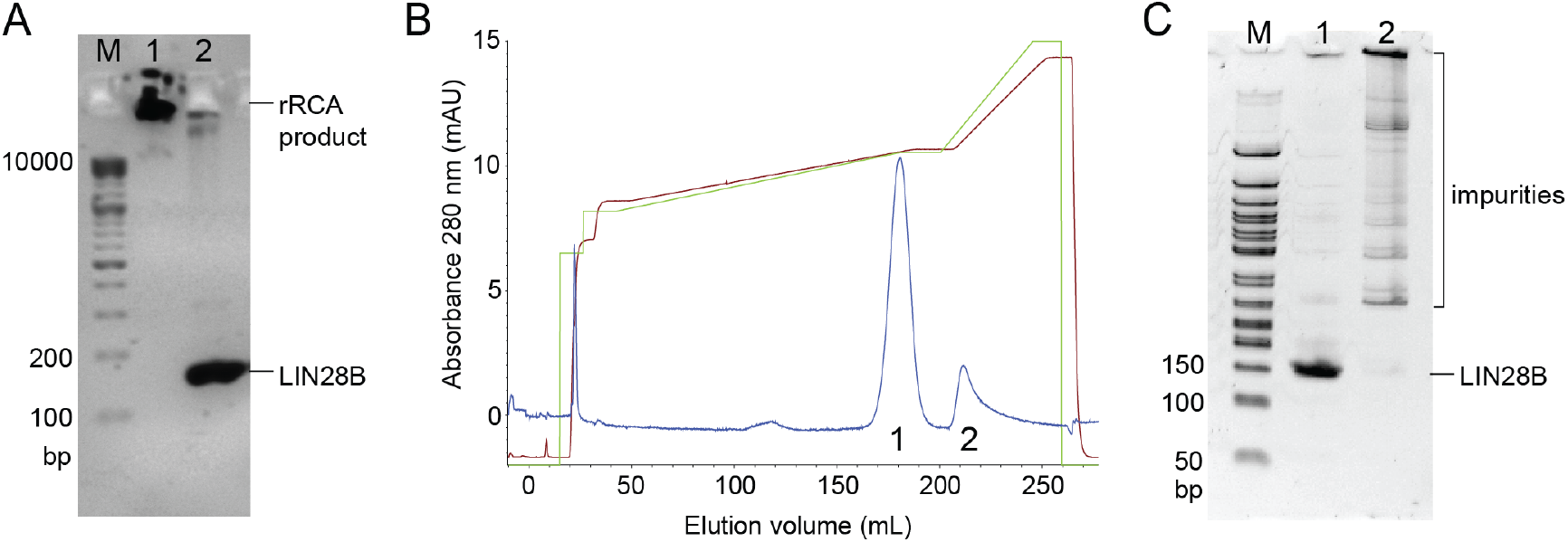
Ramified RCA, SmaI digestion and ion exchange chromatography purification. (A) Ramified RCA product (lane 1) and SmaI digestion (lane 2) (B) IEX trace of crude digested rRCA product, A280 trace in blue, applied salt gradient in green (0-100% B, 1M NaCl), measured conductivity in brown (mS). (C) 5% native PAGE gel showing the pooled fractions of peak 1 and 2, indicated in the chromatogram (panel B).

### Reconstitution of LIN28B nucleosomes and Oct4 binding

We next aimed to demonstrate that the nucleosomal DNA generated in our ramified RCA approach can be used to reconstitute functional nucleosomes. We used the LIN28B DNA from the large-scale production together with recombinantly expressed human histones to reconstitute human nucleosomes by salt-gradient dialysis, according to a previously published method (26). The reconstitution was assessed by native PAGE analysis (Figure 6A). A clear band shift was observed, indicating nucleosome formation. The efficiency of reconstitution was estimated from gel band intensities to be ~70%, which is comparable to what was previously published for LIN28B (16). We thus conclude that rRCA derived nucleosomal DNA can be used for nucleosome reconstitution in the same manner as DNA from other sources.

**Figure 6.**
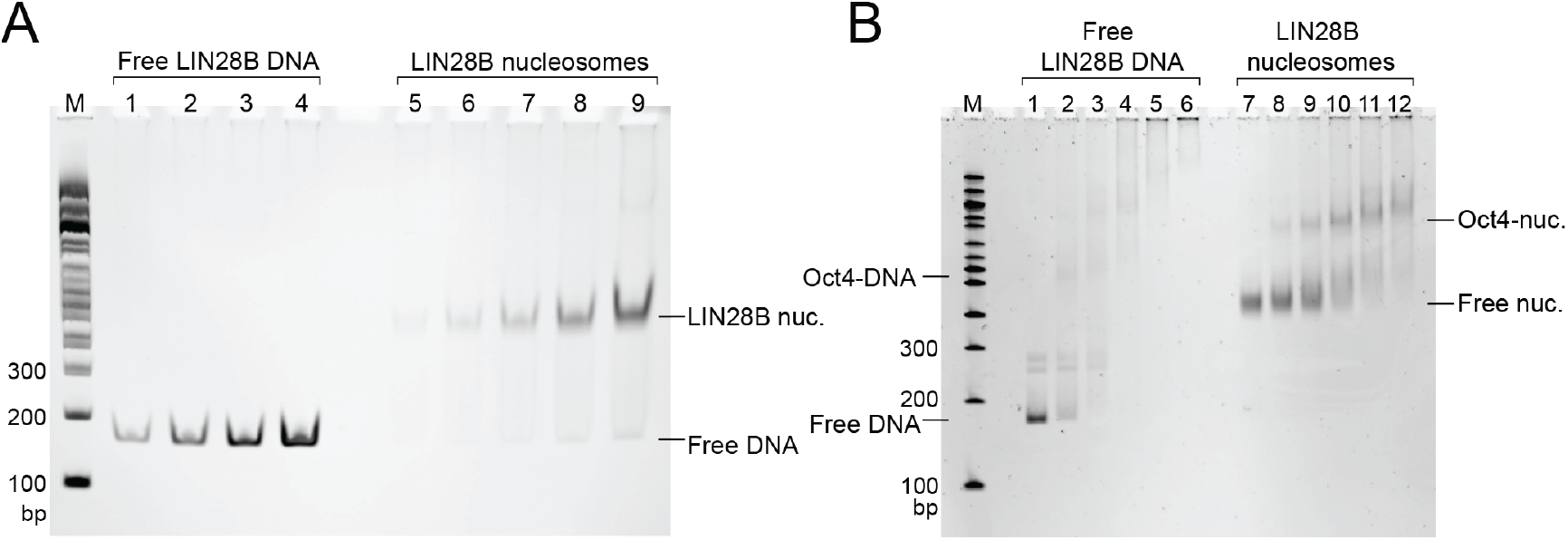
(A) Nucleosome reconstitution with rRCA-produced LIN28B DNA (162bp) and human histones. Native PAGE (6%) lanes 1-4 contain 10, 25, 50 and 100 ng of free LIN28B DNA used for quantification of the nucleosome bands. DNA was mixed with histone octamer in a 1:1.1 ratio for reconstitution. Lanes 5-9 show increasing amounts of reconstituted nucleosomes estimated to contain 7, 17, 35, 71 or 142 ng of bound DNA, respectively, which is ~70% of the input free DNA. DNA in the nucleosome has approximately a 2-fold lower signal intensity compared to free DNA. (B) EMSA of Oct4 binding to free LIN28B DNA (10 nM) and assembled LIN28B nucleosomes (25 nM). Native PAGE (5%) lanes 1-6 and lanes 7-12 contain 0, 12.5, 25, 50, 100 and 200 nM of recombinant Oct4.

To show that these nucleosomes are functional, we assayed their ability to bind pioneer transcription factor Oct4. Oct4 has been reported to associate with LIN28B nucleosomes *in vitro* in a sequence specific manner, as shown by DNase I footprinting (16). Addition of recombinant Oct4 protein to our LIN28B nucleosomes induced a clear band-shift in an EMSA experiment, indicating formation of an Oct4-nucleosome complex (Figure 6B). Both the band-shift and affinities for both free DNA and nucleosomes are in good agreement with previously published results (16). Together, these results show that the rRCA-produced LIN28B DNA has the required quality in terms of purity and sequence to enable reconstitution of native nucleosomes capable of binding pioneer transcription factor Oct4.

## DISCUSSION

We here showed that ramified RCA is an efficient and flexible alternative for the large-scale production of nucleosomal DNA in milligram quantities. Compared to plasmid and PCR-based production of nucleosomal DNA, our rRCA protocol combines the flexibility in choice of sequence from PCR with the excellent yields of a multi-repeat product from the plasmid-based method. The straightforward design procedure requires only minimal changes to the terminal ends of DNA sequence of choice in order to create the circular template needed for RCA. A crucial aspect in the design of the template is the incorporation of a restriction site to digest the long, branched product. Judicious choice of restriction enzyme allows product digestion in the same reaction volume as the rRCA reaction, boosting the time-efficiency of the method. Production, digestion and purification can be performed in two days, the whole procedure including preparation of circular templates can be completed within a week. The yield is almost 10-fold higher than in the same volume of PCR reactions. Importantly, the one-pot rRCA reaction can easily be scaled up with a linear increase in yield. We here selected a DNA sequence corresponding to a well-positioned nucleosome at the LIN28B locus. Since this sequence with 56.8% AT content has no particular features, we expect that the method can generally be applied to most DNA sequences, including larger templates for reconstitution of nucleosomal arrays on genomic DNA. Performance with highly repetitive sequences may be compromised by fidelity problems of the polymerases.

RCA requires a polymerase with extreme processivity and strand displacement, such as provided by Phi29 polymerase. Since large-scale production runs also require high amounts of enzyme, in-house production is advisable. Although this might be a barrier for labs that are not in need of continuous production of nucleosomal DNA, a 1 L culture provides Phi29 for approximately 4000 mL of rRCA and Phi29 can be stored long-term at −20°C, making even incidental use feasible.

Addition of pyrophosphatase was necessary to avoid the scavenging of Mg2+ by the formed pyrophosphate and to ensure a constant availability of Mg^2+^ throughout the reaction. This may be circumvented by performing the reaction in a dialysis bag in reaction buffer to avoid accumulation of pyrophosphate. Such a setup may improve yield by increasing buffer capacity and avoiding increases in phosphate concentration during the reaction, which could inhibit Phi29. Another potential bottleneck is the high viscosity that is observed towards the end of the rRCA reaction, since this limits the diffusion of polymerase and reactants through the solution, and thus likely reduces the reaction rate. Performing rRCA and product digestion simultaneously, preventing the formation of very long LIN28B repeats, could therefore further improve the yield. However, this requires methylation of the circular templates in combination with a methylation-sensitive restriction enzyme, to avoid digestion of the template.

The rRCA yield was improved by increasing the primer to template ratio, as well as the dNTP and Phi29 concentration. Although rRCA can provide exponential amplification of the template as demonstrated at small-scale for Bst polymerase (27), we did not observe this in our work. Analysis of the yield showed that the starting template was amplified effectively ~100-fold. The amplification here is presumably limited by the high concentration of the reactants and product leading to high viscosity. Dilution of the reaction could alleviate this, but lowered reaction rates may counteract reaching a higher yield. While further improvement in yield may be possible, we demonstrated here that our current protocol is effective and can furthermore be easily scaled. In comparison, exponential amplification in PCR is also only reached at very low template levels, thus requiring a large volume of reactions to reach milligram scale of purified product. In contrast, our rRCA protocol offers a highly practical execution without the need to prepare dozens of PCR-plates, and in particular when access to multiple PCR machines is limited. Moreover, the lack of thermocycling in rRCA leads to more efficient dNTP incorporation as a result of decreased degradation (9). Even if our protocol relies on PCR to generate the template strand, one PCR plate will generate sufficient template for a ~36 mL RCA reaction, generating ~6 mg of DNA, which would in turn be equivalent to ~120 plates of PCR.

In conclusion, we presented a rRCA-based method to allow the efficient and flexible large-scale synthesis of nucleosomal DNA sequences, any other dsDNA of comparable length. The protocol was used to produce multi-milligrams of human, genomic nucleosomal DNA with high purity. We demonstrated that rRCA-produced LIN28B DNA can be used to reconstitute stable and functional nucleosomes that are capable of binding pioneer transcription factor Oct4. Due to its ease of use and flexibility in sequence design, we believe this method is an ideal tool to produce a wide variety of nucleosomal DNA sequences to study the structure, dynamics, and interactions of nucleosomes or nucleosomal arrays, in particular of genomic nucleosomes as they occur *in vivo*.

## EXPERIMENTAL PROCEDURES

### Construction of the starting plasmid

The LIN28B template sequence was obtained as a sequence-verified gBlock^®^ gene fragment (IDT) including a 15 bp extension on both sides containing EcoRI and XbaI restriction sites for cloning into pUC19 (Table 1). Both the LIN28B gene fragment (75 ng) and the pUC19 vector (400 ng) were cut with 1U of both restriction enzymes in FastDigest buffer at 37°C for 30 min; 1U of alkaline phosphatase (FastAP, Thermo Scientific) was added to the vector restriction. Insert (75ng) and vector (25ng) were mixed and incubated overnight with 1U T4 ligase at room temperature. Ligation reaction was used to transform *E. coli* (JM109) cells and the plasmid was purified from single transformants cultured in 5 mL LB overnight and sequence-verified before further application.

### Circular template synthesis

Standard PCR reactions were performed using 2 μM of 5’-phosphorylated starting primer, 2 μM of 5’-biotinylated branching primer, 10 ng of pUC19 plasmid containing the LIN28B template sequence, 0.2 mM dNTPs and 2-3 U of home-made Pfu polymerase per 50 uL reaction. The PCR program was 3 min of initial melting at 95°C, followed by 35 cycles of 30 s at 95°C, 30 s at 50°C for annealing and 30 s at 72°C for elongation and 3 min of final polymerization at 72°C. The template strand was purified by binding the biotinylated PCR product to Streptavidin Sepharose High Performance affinity resin (GE Healthcare), purification of the beads from the reaction mixture by repeated washing with Tris buffer (10 mM Tris HCl pH 7.5, 1 mM MgCl_2_), and finally eluting the phosphorylated strand with 0.2 M NaOH. The eluent was neutralized by adding an equal volume of 0.2 M HCl and ethanol-precipitated. The template strand was circularized by heat-annealing the ligation splint and subsequently incubating this partial duplex with 200 U Taq DNA ligase (NEB) per 100 pmol of template in Taq DNA ligase buffer for 3h at 45°C. The circular template was then ethanol precipitated, reconstituted in T4 DNA ligase buffer and treated with 12.5 U Exonuclease I and 125 U Exonuclease III (Thermofisher) per 100 pmol of template at 37°C for 1 hour to remove the splint, unligated template and any other remaining single-stranded impurities. The circular templates were isopropanol precipitated and dissolved in MQ to a concentration of 1 μM before use in ramified RCA.

### Phi29 expression and assessment of activity

The gene for Phi29 DNA polymerase (Phi29DNAP) was amplified by PCR from a stock of Phi29 phages using primers 5’ACCATGGATCC<u>CATATG</u>CCGAGAAAGATGTATAG3’ and 5’ACCAT<u>GAATTC</u>TCGAGTTATTTGATTGTGAATGTG3 (restriction sites underlined) and cloned into the NdeI and EcoRI sites of pET28a (Novagen) introducing a N-terminal His6-tag. *E. coli* BL21(DE3) was transformed with plasmid pPhi29DNAP and cultivated in LB medium under kanamycin selection at 37°C. Three baffled Erlenmeyer flasks containing 1 liter media were inoculated with 25 mL overnight culture and grown at 37°C with shaking at 225 rpm to an OD600 of ~0.8. IPTG was added to 1 mM and cultivation continued for another 3 hours. Cells were harvested by centrifugation at 5000 rpm, 4°C, 10 minutes in a Beckman JA-10 rotor. Alternatively, Phi29 polymerase was expressed in *E. coli* Rosetta2 cells in 250 mL of auto-induction medium (ZYM-5052) at 27.5°C overnight (28). Cells were resuspended in ice-cold lysis buffer (25 mM Tris-HCl (pH7.5), 0.5% Tween-20, 0.5% Nonidet P-40 substitute, 5% glycerol, 20 μg/mL PMSF, 5 mM β-mercaptoethanol, 1 mM EDTA and 100 μg/mL lysozyme) and sonicated 10 cycles, each cycle consisting of 30 seconds at 10 micron amplitude and 1 minute chill on ice. MgCl_2_ was added to 2.5 mM, CaCl_2_ to 0.5 mM and DNaseI to 2 U/mL and the lysate was stirred for 10 minutes at room temperature. Imidazol was added to 10 mM, EDTA to 3 mM and NaCl to 300 mM and the lysate was then centrifuged for 30 minutes at 15000 rpm, 4°C in a Beckman JA-20 rotor. The supernatant was loaded onto a HisTrap HP Sepharose column (5 mL, GE Healthcare) pre-equilibrated in column buffer (25 mM Tris-HCl (pH7.5), 0.5% Tween-20, 0.5% Nonidet P-40 substitute, 5% glycerol, 5 mM β-mercaptoethanol, 10 mM imidazol and 300 mM NaCl). The column was washed with 25 mL of column buffer containing 25 mM imidazol and Phi29 DNAP was eluted in fractions of 1 mL with column buffer containing 300 mM imidazol. Fractions containing Phi29DNAP were pooled and added to 50 mL of nuclease treatment buffer (25 mM Tris-HCl (pH7.5), 0.5% Tween-20, 0.5% Nonidet P-40 substitute, 5% glycerol, 5 mM β-mercaptoethanol, 2.5 mM MgCl_2_, 0.5 mM CaCl_2_, 100 U DNaseI, 200 U exonuclease I and 200 U RNaseI) and stirred for 10 minutes at room temperature. EDTA was added to 3 mM and NaCl to 300 mM and the HisTrap HP Sepharose column purification was repeated on a freshly regenerated column. Protein containing fractions were pooled and dialyzed overnight against 1 liter of dialysis buffer (50 mM Tris-HCl (pH7.5), 100 mM NaCl, 1 mM DTT, 0.1 mM EDTA, 0.5% Tween-20, 0.5% Nonidet P-40 substitute and 50% glycerol) at 4°C. The Phi29 DNAP isolate was aliquoted and stored at −20°C. Activity of the isolate was determined by comparing product DNA yields relative to a commercial isolate (Epicentre).

### Ramified RCA and digestion

The LIN28B circular template (20 nM) was amplified in a large-scale ramified RCA reaction in 1x CutSmart^®^ buffer (NEB) with 200 U of home-made Phi29 polymerase and 0.5 U inorganic pyrophosphatase (NEB) per mL of reaction, 0.5 mM dNTPs, 2 μM of starting and branching primer and 4mM DTT. Circular template and primer were heat-annealed in CutSmart buffer before adding dNTPs and the enzymes. The rRCA reaction was incubated at 30°C for 5h, followed by the addition of 1000 U SmaI (NEB) per mL of reaction and further incubation at 25 °C overnight. The resulting double stranded 162bp LIN28B DNA was purified by anion exchange chromatography and ethanol precipitated.

### Nucleosome reconstitution

rRCA-produced LIN28B DNA was mixed with human purified histone octamers at a 1:1.1 DNA:octamer molar ratio in the presence of 2M NaCl and nucleosomes were reconstituted by salt gradient dialysis. The nucleosomes were analyzed on 6% TBE gel (Novex) in 1X TBE at 90 V for 1 hr and visualized by Ethidium Bromide staining. The nucleosome concentration was calculated by quantifying the intensities of nucleosome bands, using free LIN28B DNA as the standard. The densitometric analysis of band intensities was performed using Multi-Gauge software (Fujifilm Science lab).

### Mobility shift assay with Oct4

Full-length human Oct4 fused to an N-terminal 6X histidine tag with a thrombin cleavage site was expressed from a pET28b bacterial expression plasmid in *E. Coli* Rosetta (DE3) pLysS cells. The recombinant protein was purified under denaturing conditions over a His-SpinTrap column (GE Healthcare), desalted using a PD SpinTrap G-25 column (GE Healthcare) and concentrated using an Amicon Ultra-0.5 device (MW cut-off 10 kDa). EMSA was performed with increasing amounts of Oct4 to free LIN28B-DNA and LIN28B-nucleosomes. The free LIN28B-DNA (approx. 10 nM) and LIN28B-nucleosomes (approx. 25 nM) were incubated with recombinant Oct4 protein (12.5, 25, 50, 100, 200 nM) in DNA-binding buffer (10 mM Tris-HCl (pH 7.5), 1 mM MgCl_2_, 10 μM ZnCl_2_, 1 mM DTT, 10 mM KCl, 0.5 mg/ml BSA, 5% glycerol) at room temperature for 60 min. Free and Oct4 bound DNA were separated on 5% non-denaturing polyacrylamide gel run in 0.5X TBE at 90V for 4 hrs. The gel was stained with ethidium bromide and visualized using SynGene G:Box.

## Acknowledgement

We thank dr. Gert Folkers for stimulating discussions and helpful suggestions.

## FOOTNOTES

This work was supported by a VIDI grant from the Dutch Science Foundation NWO to HvI (NWO-CW VIDI 723.013.010). AS was supported by an MRC career development award (MR/N024028/1). IG was supported by the BBSRC-EASBIO Ph.D training program (BB/M010996/1).

## The abbreviations used are

rRCA: ramified rolling circle amplification;
PCR: polymerase chain reaction;
dNTPs: deoxyribose nucleoside triphosphates

